# Deletion of the short N-terminal extension in OCP reveals the main site for the FRP binding

**DOI:** 10.1101/116491

**Authors:** Nikolai N. Sluchanko, Yury B. Slonimskiy, Marcus Moldenhauer, Thomas Friedrich, Eugene G. Maksimov

**Affiliations:** A.N. Bach Institute of Biochemistry, Federal Research Center of Biotechnology of the Russian Academy of Sciences, 119071 Moscow, Russian Federation; M.V. Lomonosov Moscow State University, Department of Biophysics, Faculty of Biology, 119991 Moscow, Russian Federation; M.V. Lomonosov Moscow State University, Department of Biochemistry, Faculty of Biology, 119991 Moscow, Russian Federation; Technical University of Berlin, Institute of Chemistry PC 14, Straβe des 17. Juni 135, D-10623 Berlin, Germany

**Keywords:** Orange carotenoid protein, fluorescence recovery protein, phycobilisome fluorescence, protein-protein interactions, photoactive proteins, photoprotection

## Abstract

The photoactive Orange Carotenoid Protein (OCP) plays a central role in cyanobacterial photoprotection. Photoconversion entails significant structural rearrangements in OCP required for its binding to the phycobilisome to induce excitation energy dissipation, whereas the fluorescence recovery protein (FRP) is required for OCP detachment and restoration of phycobilisome fluorescence. Although key to understanding the whole reversible mechanism of photoprotection, the FRP binding site on OCP has been representing challenge since the discovery of FRP in 2010 and is currently unknown. OCP comprises two structural domains organized into a compact basic orange form due to specific protein-chromophore and inter-domain protein-protein interactions and interacts with FRP tightly only when photoactivated. As an important stabilizing element in the orange OCP, the short αA-helix within the N-terminal extension (NTE) binds to OCP’s C-terminal domain (CTD), but unfolds upon photoactivation and interferes with phycobilisome binding. By using an alloy of biochemical and biophysical techniques, here we demonstrate that the NTE shares specific structural and functional similarities with FRP and discover the main site of OCP-FRP interactions in the OCP-CTD.

## 1. Introduction

As any extreme, Life’s virtually insatiable want for photons needs taming by mechanisms of temperance. To permit growth even in the shade, photosynthetic organisms developed meticulous light-harvesting or antennae complexes to optimize capture efficiency for feasting on essentially every photon. However, exposure to intense sunlight potentially entails fatal photooxidative damage of the photosynthesis machinery, especially the reaction centers. Cyanobacteria have evolved a light-regulated system of photoprotection that controls the flow of excitation energy from their soluble light-harvesting complexes, the phycobilisomes (PBs), to the reaction centers of the photosystems [1]. Pivotal for this mechanism is the photoactive 34.7 kDa Orange Carotenoid Protein (OCP) [2-4], which harbors a special xanthophyll as active pigment and undergoes phototransformation from an orange state (OCP^O^) to the active red state (OCP^R^) (see [5] for the review). This photoconversion is accompanied by substantial conformational changes, which permit tight interaction of the OCP^R^ state with PBs to effectively quench their fluorescence, thereby shielding the reaction centers from excessively absorbed excitation energy [4, 6-8]. While the OCP^O^ to OCP^R^ transition requires absorption of a photon, the back conversion occurs spontaneously in the dark, but *in vivo,* the process of termination of PBs fluorescence quenching invokes the action of the Fluorescence Recovery Protein (FRP) [9-11], which stimulates detachment of OCP from the PBs and speeds up OCP^R^ to OCP^O^ back-conversion [12, 13]. Quenching of PBs fluorescence by light intensity-dependent formation of OCP^R^ and tight OCP^R^/PBs interaction, antagonized by the action of FRP, is the cyanobacterial variant of a more general photoprotective mechanism termed non-photochemical quenching (NPQ), which manifests as quenching of chlorophyll fluorescence that is active in many photosynthetic organisms including higher plants and algae [14-16].

According to the available crystal structures, OCP can be subdivided into two domains of about equal molecular weight, a 17.7 kDa predominantly α-helical unique N-terminal domain (NTD) and an 17 kDa mixed α--helical/pβ-sheet-containing C-terminal domain (CTD), the latter bearing similarity to the superfamily of nuclear transport factor-2 (NTF-2)-like protein domains (PFAM 02136) that is widely distributed among pro- and eukaryotes [17-20]. Both domains encapsulate the carotenoid in a central cavity that near symmetrically stretches into NTD and CTD, leaving only about 4 % of the carotenoid surface exposed to the aqueous environment. The crucial role of the 4-(or 4’-)keto oxygen at the terminal β-ring(s) is underlined by the fact that in the crystal structures of the basic OCP^O^ state, the only specific carotenoid-protein interactions are two H-bonds formed in CTD between the N-H imino group of Trp-288 and the hydroxyl of Tyr-201 with the 4-(4’-)keto oxygen of the xanthophyll. Besides these protein/chromophore interactions, the compact OCP^O^ state is stabilized by inter-domain interactions via the large NTD/CTD interface, including the Arg-155/Glu-244 salt bridge, and, importantly, hydrophobic interactions between the residues from α-A-helix within the about 20 amino acids-long N-terminal extension (NTE) of NTD and a specific surface area involving several pβ-sheets of CTD [21-23]. The structure of the red signaling state OCP^R^ is still elusive though being a prerequisite for understanding the interactions of the active form with PBs and FRP. It is commonly accepted that photon absorption causes carotenoid isomerization, and the subsequent breaking of the critical H-bonds initiates NTD-CTD separation as well as detachment and unfolding of the αA-helix [22-26]. This results in substantially increased volume of OCP^R^ with characteristics of a molten globule [25, 27]. Comparison of the crystal structures of OCP and its isolated NTD, which is also known as Red Carotenoid Protein (RCP), suggested that, upon photoconversion, the carotenoid cofactor completely leaves CTD and shifts by 12 AÅ into NTD [20], a process proven to occur dynamically in full-length OCP by means of distance measurements based on Förster resonance energy transfer [26].

Although the structure of FRP is known, the mechanism of its interaction with the OCP remains far from being clear. We have recently introduced the purple OCP^W288A^ variant [27] as a fully competent analog of the OCP^R^ signaling state, which does not require photoactivation, constitutively quenches PBs fluorescence and binds FRP with micromolar affinity, whereas FRP binding to OCP^O^ was about 10-fold weaker [28]. High-affinity FRP binding was also shown for the OCP apoprotein, which also exhibits substantially increased volume suggesting that FRP interacts with essentially any structurally extended form of OCP [29]. Thus, FRP may serve as a scaffold to reverse domain separation and allow for re-formation of the basal OCP^O^ state as well as serving in proper chromophore insertion during maturation of the OCP protein. FRP/OCP interaction occurs with 1:1 stoichiometry implying the dissociation of FRP dimers upon binding to OCP [28, 29]. The regions on FRP and OCP involved in these physiological interactions are still controversial. It has been shown that FRP predominantly binds to the isolated CTD of OCP [11, 21, 29] and recent mutagenesis studies have shown that several amino acids (especially F299) within the interface on CTD with the N-terminal αA-helix modify photoactivation/back-conversion kinetics and FRP interactions in the course of PBs fluorescence quenching [13]. Interference of the αA-segment with PBs binding has also been inferred [21]. However, the FRP segments involved in OCP interaction are less well known. According to the crystal structure [11], FRP harbors mainly α-helical structure elements but may assume two largely different configurations, even with different oligomeric state, suggesting that large conformational changes might occur, especially bending between α1- and α2-helix. Molecular docking paired with mutagenesis proposed that a segment on FRP-α2 may bind to a region on the OCP-CTD overlapping with the binding site of OCP-αA [11], however, this docking was performed with dimeric FRP and entailed substantial clashes with the OCP-NTD.

In this work, we have tested the hypothesis of competitive binding of FRP and the αA-helix of the OCP-NTE to one common site defined by αA-binding region of the OCP^O^ structure. On the basis of amino acid sequence similarities of the α-helical N-terminal segment of FRP (α1) and OCP-αA, we propose the primary (active) binding region of FRP as overlapping with that for the NTE. We show that FRP binds even with sub-micromolar affinity and 1:1 stoichiometry to an OCP variant lacking the NTE (ΔNTE), stays bound during photoactivation of the latter, and greatly accelerates red-to-orange back-conversion of the ΔNTE protein, with even 5-fold higher rate than in the case of wildtype OCP (OCP^WT^). All in all, these data support the idea that the binding site of FRP and OCP-αA on the OCP-CTD are identical and mutual binding occurs in a competitive fashion.

## 2. Results and Discussion

### 2.1. Obtaining and characterization of the two distinct fractions of ΔNTE

An affinity-purified ΔNTE protein sample expressed in ECN/CAN-producing *E. coli* cells appeared red (Fig. S1A), in line with the observations by Kirilovsky and co-workers [21]. Surprisingly, upon separation of the sample by SEC, two fractions having substantially different elution volume and color were obtained - the earlier eluted *purple* and later eluted *orange* (Fig. S1A). By analogy with the color of the earlier described purple OCP mutant [27, 28] and the wildtype orange OCP (OCP^WT^), we termed these fractions ΔNTE(P) and ΔNTE(O), respectively. Expanding the analogy, in line with our earlier data [21], on the analytical SEC, we observed that ΔNTE(P) showed a characteristic concentration-dependent dimerization (Fig. S1B,C), in contrast to ΔNTE(O) that showed only a single symmetrical peak (~27.5 kDa) irrespective of protein concentration. Importantly, the ΔNTE(P) fraction contained a large amount of apoprotein, which, in line with our earlier observations [30], also displayed concentration-dependent self-association and, therefore, co-eluted with ΔNTE(P) leaving the ΔNTE(O) fraction almost free from any apoprotein.

**Figure 1.**
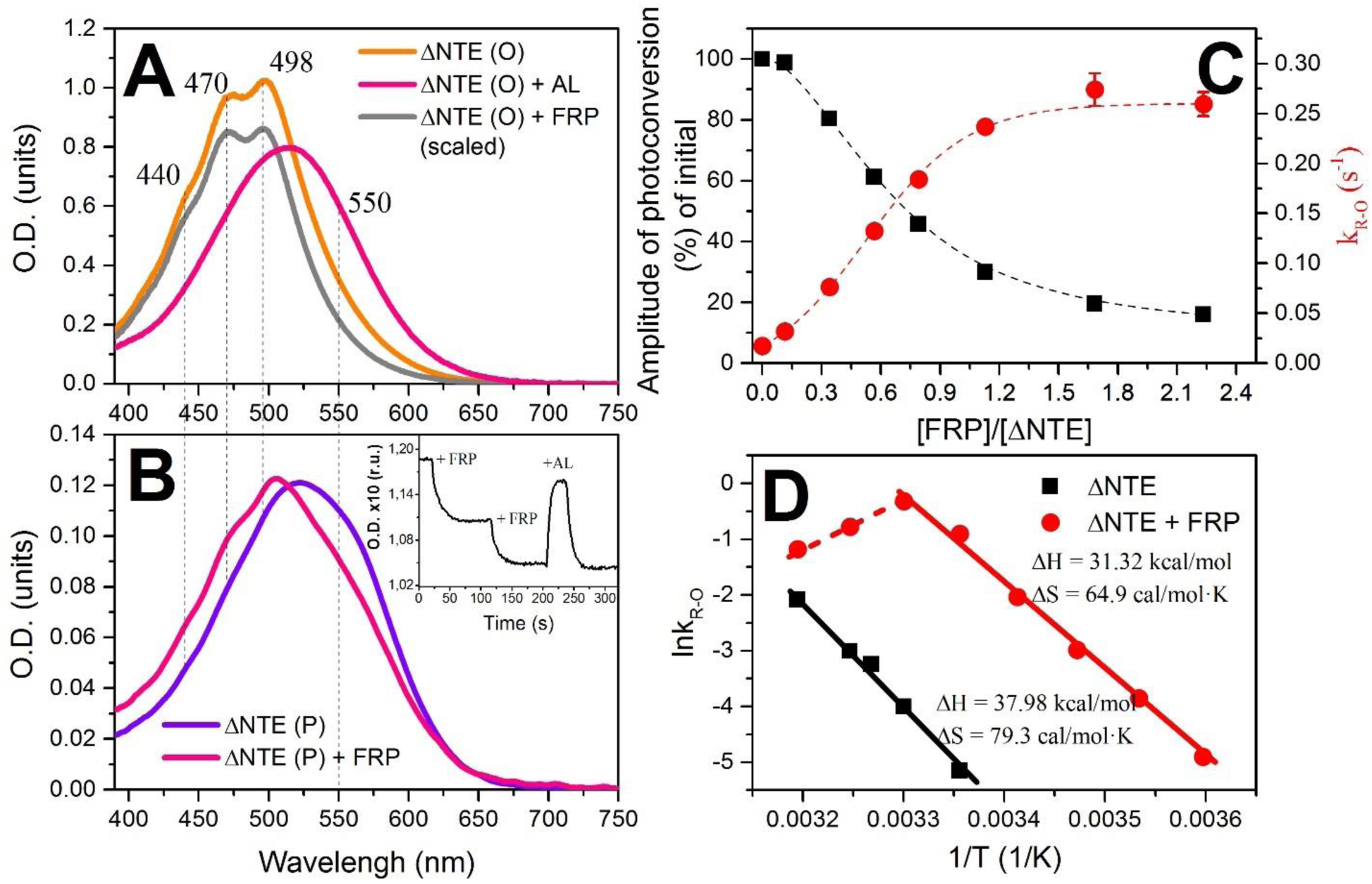
Absorption spectra of the (O)range and (P)urple species of the OCP ΔNTE protein before and after addition of FRP in excess. In (**A**), the spectrum of ΔNTE after addition of FRP (gray) is down-scaled for better presentation. In (**B**), the inset indicates a time-course of optical density (O.D.) changes at 550 nm upon addition of two FRP aliquots and following illumination by actinic light (AL). Experiments were conducted at 25 °C with constant stirring of the samples. (**C**) Dependency of the photoconversion amplitude (black squares, measured as maximal change in the O.D. at 550 nm upon illumination by 200 mW of actinic light) and the red to orange relaxation (R→O) rates (red circles) on concentration of FRP added to 2.5 μM solution of ΔNTE(O). Experiments were conducted at 25 °C. (**D**) Arrhenius plot of R→O relaxation rates in the absence (black squares) and in the presence of FRP (red circles). Experiments were conducted in the range of temperatures from 5 to 40 °C.

### 2.2. Interactions of ΔNTE and FRP as revealed by absorption spectroscopy

The absorption spectrum of the ΔNTE(O) fraction was similar to the one of ECN-coordinating wildtype OCP, with an increase of absorption in the red part of the spectrum, resulting in higher intensity of the 498 nm band compared to the intensity of another vibronic peak at 470 nm *(Fig. 1A).* This observation is in a good agreement with a previously reported presence of about 30% red OCP form in the sample of Δ 2-20 and Δ 2-15 OCP mutants [21]. Equilibrium between orange and red states was sensitive to temperature (data not shown) with less red form at high temperatures. Illumination of ΔNTE(O) by actinic light caused changes of absorption characteristic for OCP resulting in formation of the red form *(Fig. 1A).* Surprisingly, addition of FRP to the solution of ΔNTE(O) caused changes of the absorption spectrum, implying that FRP is able to interact with ΔNTE(O) in its orange state *(Fig. 1A),* which is not characteristic for OCP^WT^ (or occurs with extremely low affinity [28]). The resulting spectrum *(Fig. 1A* gray line) was identical to that of OCP^WT^. This suggests that FRP causes fine-tuning of the protein-carotenoid and protein-protein interactions in OCP, and – in this case – directly aids in formation and stabilization of the orange state, consistent with our earlier observations [28].

The most significant changes of absorption spectra were observed upon addition of FRP to the purple ΔNTE(P). This procedure not only allowed to obtain the sample with characteristic signatures of the orange form, but also to *restore the photoactivity* of the sample *(Fig. 1B inset).* Thus, both forms of ΔNTE are able to interact with FRP in the dark.

In order to characterize the interaction of the ΔNTE variant of OCP with FRP, the rates of photocyclic transitions of ΔNTE(O) were studied by monitoring the changes of absorption. In principle, we could also have used the ΔNTE(P) sample with restored photoactivity, however, in this case, the amplitude of photoconversion was extremely small at low FRP concentrations, which would affect the accuracy of rate constants determination. Due to this fact and taking into account the substantial contamination of ΔNTE(P) by the apoprotein, we further present the results obtained on ΔNTE(O) only.

It should be noted that the rate of the R→O conversion for ΔNTE appeared (slightly lower but) comparable to the one of OCP^WT^, indicating that the presence of NTE (including the αA-helix) is not indispensable for re-association of CTD and NTD and formation of the orange state. Addition of FRP to ΔNTE gradually increased the rate of the R→O conversion by more than an order of magnitude *(Fig. 1C),* and saturated at about 1:1 molar ratio. The increase in the rate of R→O conversion leads to a reduction of the amplitude of O.D. changes upon photoconversion, consistent with our previous results [28] and a kinetic model [25]. Surprisingly, the rate of the R→O conversion in the presence of FRP was almost 5 times higher for the OCP ΔNTE variant compared to the corresponding rates of OCP^WT^ relaxation at exactly the same FRP/OCP concentration ratio determined previously [28]. This fact not only proves that the NTE is not necessary for the R→O conversion of OCP, but also points out that this structural element may need to sterically compete with FRP for the specific site on OCP. Obviously, the NTE also delays closure of the protein and formation of the orange state, as seen from the appreciable reduction of the R→O relaxation rates, which could be crucial since maintaining OCP in the active state determines quenching of phycobilisome fluorescence. Analysis of temperature dependencies of the R→O rates for the ΔNTE(O) variant in the absence and in the presence of FRP *(Fig. 1D)* shows that formation of the tentative ΔNTE/FRP complex reduces both, enthalpy and entropy, of the transition by ~18 %, i.e. slightly more profound compared to the corresponding characteristics of the OCP^wt^/FRP interaction [28]. It should be noted that at high temperatures above 30 °C we observed strongly non-linear behavior of R→O rates in the presence of FRP due to the appearance of a slow component in the kinetics of R→O transition, which is probably related to reduction of FRP assistance. Previously, we observed similar reduction of the rates at high temperatures for OCP^WT^ in the presence of FRP [28], however in the absence of NTE this effect is much more pronounced.

### 2.3. Direct interaction of the ΔNTE variant with FRP

Recently, we showed that the analog of the red OCP form, the purple OCP^W288A^ mutant, interacts with FRP at a 1:1 stoichiometry and micromolar apparent affinity, whereas the interaction with the basic orange OCP^WT^ is about 10 times weaker [28]. We hypothesized whether the absence of NTE opens the possibility for a tight physical interaction between OCP and FRP and checked this hypothesis by analytical SEC. Indeed, while the individual FRP and ΔNTE(O) peaks had almost exactly the same position (~27-28 kDa), by following carotenoid-specific absorbance at 500 nm (i.e., detecting exclusively holo-ΔNTE(O) in the bound and unbound states), we could analyze the elution profile changes associated with complex formation. Increasing concentrations of FRP led to a gradual migration of the ΔNTE(O) peak from ~13.0 min to ~12.2 min retention time *(Fig. 2A),* and this corresponded to an apparent molecular weight increase from 27.5 to ~40 kDa, unequivocally indicating equimolar complex formation between ΔNTE(O) and FRP upon FRP monomerization (27.5 + 14 kDa). Approximation of the binding curves obtained using a series of FRP concentrations confirmed the 1:1 stoichiometry and resulted in a submicromolar apparent K_D_ *(Fig. 2B)*. Intriguingly, this affinity is the highest between FRP and any OCP variant observed so far. From a kinetic point of view, taking into account the substantially larger apparent K_D_ of 2-3 μM determined recently for the FRP interaction with the red OCP form analog OCP^W288A^ [28], the difference in the aforementioned dissociation constants is exactly what one would expect for competitive binding of FRP and the NTE to a single site on the OCP-CTD. In the presence of the competing NTE (as in OCP^W288A^) the apparent affinity of FRP binding is appreciably lower than in its absence. The different K_D_ values of orange and red analog forms may also suggest that the FRP-binding site on OCP might comprise more than one region, but one of these certainly overlaps with the NTE-binding site. In either of these scenarios, our results support the scaffold-like role for FRP relative to OCP. In principle, we could also detect FRP interaction with ΔNTE(P), which showed saturation (Fig. S2A), however, in this case it was difficult to determine the binding parameters quantitatively due to the inevitable apoprotein contamination of the ΔNTE(P) sample (*Fig. S2B*).

**Figure 2.**
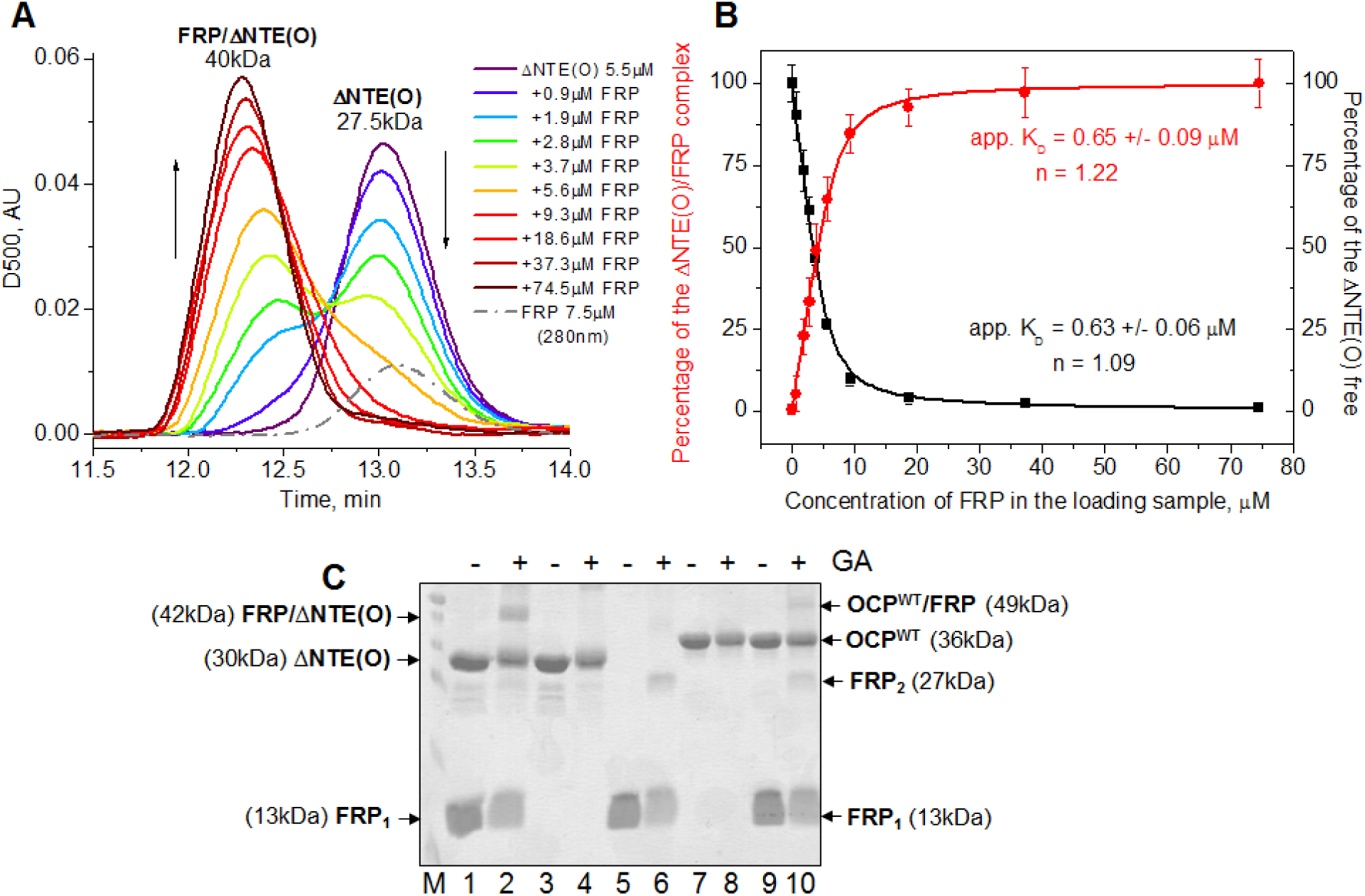
Interaction of the ΔNTE(O) variant of OCP with FRP studied by SEC and chemical crosslinking with glutaraldehyde (GA). (**A**) SEC profiles for ΔNTE(O), FRP, or their mixtures at increasing FRP concentration (in the range 0-74.5 μM) obtained using a Superdex 200 Increase 10/300 column at a flow rate of 1.2 mL/min. Positions of ΔNTE(O), its complex with FRP and their apparent molecular weights are indicated. (**B**) Binding curves reflecting changes in the amplitudes of either disappearing peak of free ΔNTE(O) (black) or appearing complex peak (red) seen in panel (**A**). Approximation was done using a quadratic binding equation (see Materials and Methods and [28]), which revealed almost stoichiometric binding and apparent K_D_ values. (**C**) Chemical crosslinking of the ΔNTE(O)/FRP mixture (lanes 1, 2), individual ΔNTE(O) (lanes 3, 4), individual FRP (lanes 5, 6), individual OCP^WT^ (lanes 7, 8), or its mixture with FRP (lanes 9, 10). Uncrosslinked (odd lanes) and crosslinked by 0.01% GA (even lanes) samples were analyzed. Assignments of protein bands and their apparent molecular weights are indicated by arrows. M – molecular weight markers (from bottom to top: 10, 15, 25, 35, 40, 50 kDa). FRP_1_ and FRP_2_ correspond to FRP monomers and dimers, respectively. Each experiment was done at least three times.

Significantly, we could confirm the binding stoichiometry and the size of the ΔNTE(O)/FRP complex by chemical crosslinking with glutaraldehyde (GA). Under conditions used, FRP showed the formation of crosslinked dimers, whereas neither OCP^WT^ nor ΔNTE(O) oligomerized in the dark-adapted form *(Fig. 2C).* Of note, in SDS-PAGE, ΔNTE runs faster than OCP^WT^ due to lower molecular weight. At the same time, we could detect formation of the ~42 kDa band upon cross-linking the ΔNTE(O)/FRP mixture *(Fig. 2C,* lane 2), and the absence of this band in other samples indicates that it corresponds to the 1:1 complex between the two proteins (13+30 kDa). We could confirm the presence of 1:1 crosslinked complex also in the OCP^WT^/FRP mixture, however, its intensity was significantly lower *(Fig. 2C*, compare lanes 10 and 2) in line with the lower affinity of interaction [28], additionally confirming the tight, specific, and light-independent interaction between FRP and the OCP variant devoid of NTE.

### 2.4. The pre-formed ΔNTE/FRP complex is preserved upon intense blue-light illumination

Although FRP binds with high affinity to the dark-adapted OCP variant lacking the NTE, the important question is whether this interaction is possible upon illumination and tentative photoactivation of ΔNTE. As expected, illumination by a blue LED during separation on a SEC column resulted in shifts towards larger particle sizes of both OCP^WT^ and ΔNTE(O) species. Moreover, the amplitudes of the shifted peaks registered at 500 nm absorbance decreased (*Fig. S3A*), implying photoactivation of both OCP variants and formation of their red forms with the red-shifted absorption spectrum and larger size [27, 28]. Since the orange forms of both variants display low concentration-dependent dimerization [28], the difference in the elution time (*Fig. S3* A) is associated with the deletion in the ΔNTE variant.

Importantly, as in Fig. 3A, addition of FRP in the dark (Fig. S3B) resulted in a dramatic shift of the ΔNTE peak, reflecting formation of their complex. However, illumination of the samples during the chromatography run with blue-LED at maximum power (900 mW) led to only a barely detectable shift of the complex peak toward larger sizes (Fig. S3B). This indicates that (i) FRP stays tightly bound to ΔNTE even upon ΔNTE photoactivation and (ii) that the R→O relaxation in such a system is so fast, that we are able to detect mostly the already relaxed O-like compact state of ΔNTE. According to absorption measurements, FRP binding does not preclude ΔNTE(O) from photoactivation-associated “redding”, although reduces it by ~80 % (see *Fig. 1C).* According to Fig. S3B, photoactivation of ΔNTE(O), in turn, does not exclude FRP binding, and the latter remains bound to CTD even upon NTE and NTD separation and, thus, efficiently stimulates R→O conversion. This scenario resembles the well-known *rodeo,* but on a molecular level, as FRP tames its furious horse (OCP^R^) by making it quiet and calm (OCP^O^). Our principal finding implies that, in normal OCP^WT^, NTE outcompetes FRP from the complex with OCP which has already been converted back to its basic orange state and permits further FRP action on the OCP molecules.

**Figure 3.**
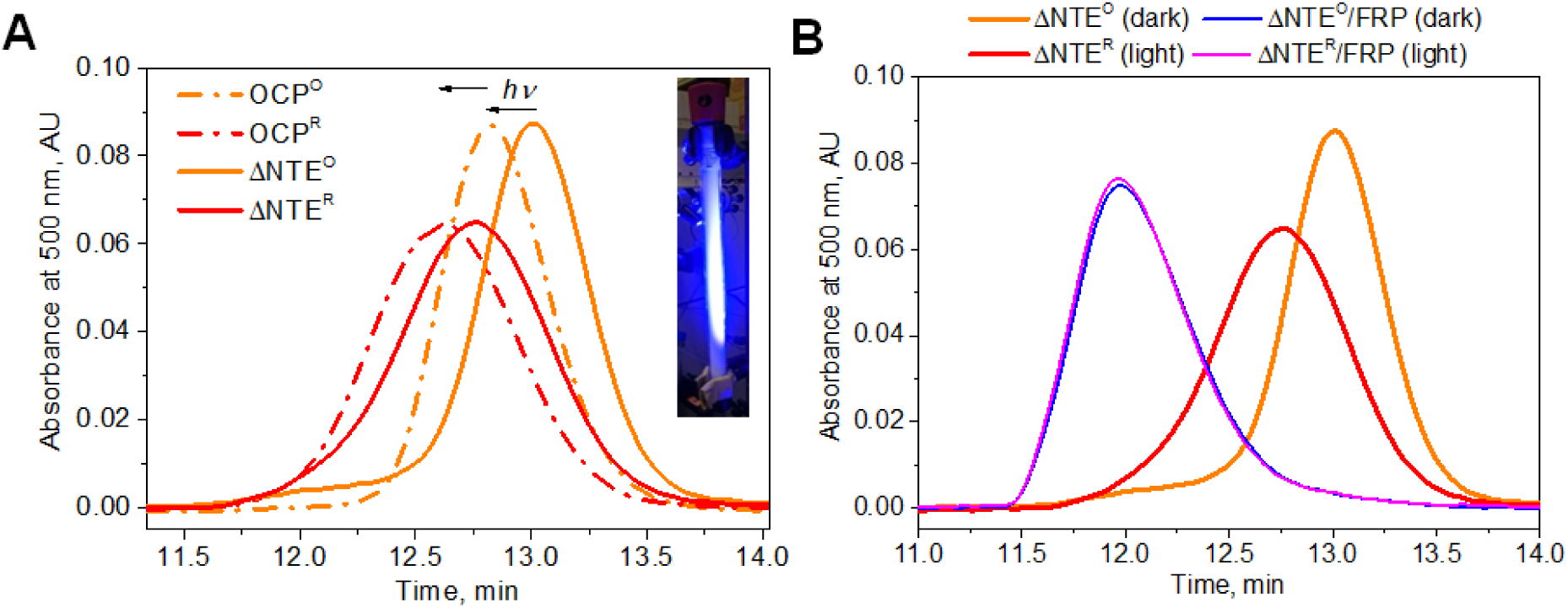
The effect of blue light illumination on the SEC elution profiles of ΔNTE and OCP^WT^ (**A**) Photoactivation of OCP and the ΔNTE variant results in a decrease of absorption at 500 nm and an increase of the size of the particles, in a similar fashion. The insert shows the experimental setup with the Superdex 200 Increase column illuminated by blue-LED in the course of sample separation. (**B**) The effect of high light illumination on the elution profile of ΔNTE in the absence or in the presence of FRP. The experiment was repeated twice with similar results.

## 3. Conclusions

In Figure 4, we propose a structural and kinetic model of the mutual competition of FRP and the NTE of OCP for binding on the OCP-CTD. In the ΔNTE variant of OCP, the orange state is destabilized due to the absence of αA-helix, therefore, the sample represents a mixture of the orange-like (major) and red-like (minor) states. Addition of initially dimeric FRP to the ΔNTE protein results in FRP monomerization (not included in the scheme of *Fig. 4D* see [28]). Binding of an FRP monomer to the site on the CTD of ΔNTE in effect promotes closure of the OCP domains and causes stabilization of the carotenoid position making the color of the sample identical to OCP^WT^ in its orange state. In this binding mode, FRP recognizes the main (primary) site for the attachment in the CTD of OCP, which is occupied by the αA-helix of the NTE in wildtype OCP^O^ but is freely accessible in the OCP^R^ state. Upon binding to the main site on OCP-CTD, FRP monomerization occurs, causing exposition of groups capable of specific protein-protein interactions with both structural domains of OCP. Due to the high affinity of ΔNTE-OCP^O^ towards FRP and absence of NTE, the complex does not dissociate in the dark *(Fig 3)*. Under high intensity of blue-green light, isomerization of the carotenoid initiates ΔNTE-OCP^O^ photoconversion into the active red form ΔNTE-OCP^R^, which results in breaking of critical H-bonds including the one between Trp-288 and the keto-group of the carotenoid [25, 27, 30, 31]. However, since the main site of OCP/FRP interactions is already occupied by FRP in the orange ΔNTE-OCP^O^/FRP complex, FRP immediately provides a tentative *scaffold-like* assistance, which allows the ΔNTE protein to orient CTD and NTD properly (facing each other) and to rapidly recover into its initial conformational state due to establishment of previously disrupted bonds (including the Arg-155/Glu-244 salt bridge), and thus promotes compaction of the protein and an overall decrease in entropy and enthalpy of the complex *(Fig. 1D).* As shown in Fig. 4C, the N-terminal sequence AETQSAHALFR belonging to α1-helix of FRP is homologous and structurally match to the sequence FTIDSARGIFP in αA-helix of OCP’s NTE (see alignment in lower panel of Fig. 4). This FRP binding model is consistent with several established hydrophobic contacts between highly conserved residues of OCP, including the experimentally validated F299 [13], and FRP, however, is somewhat different from the earlier predicted “active site” of FRP (residues 50-60) [11]. Nevertheless, this “active site”, which was shown to be important for the accelerating effect of FRP on R→O conversion of OCP, may be involved in FRP dimerization or in formation of the tentative second site of FRP binding on OCP-NTD, necessary for the closure of OCP during the R→O transition. The proposed scaffold-like action of FRP can benefit from the ability of this small α-helical protein to flexibly adopt different structures as evidenced by two distinct conformations in the crystal structure (PDB entry 4JDX) and its ability to monomerize upon OCP binding [11, 28]. The existence of a secondary FRP binding site on the OCP-NTD is indirectly supported by the difference between the apparent dissociation constants measured for FRP binding to the ΔNTE variant of OCP (<1 μM) and to the red signaling state analog OCP^W288A^ (2-3 μM [28]). However, location of the secondary site is currently unclear, and further studies, including structural biology work, are clearly needed to validate the proposed OCP/FRP interaction mode.

**Figure 4.**
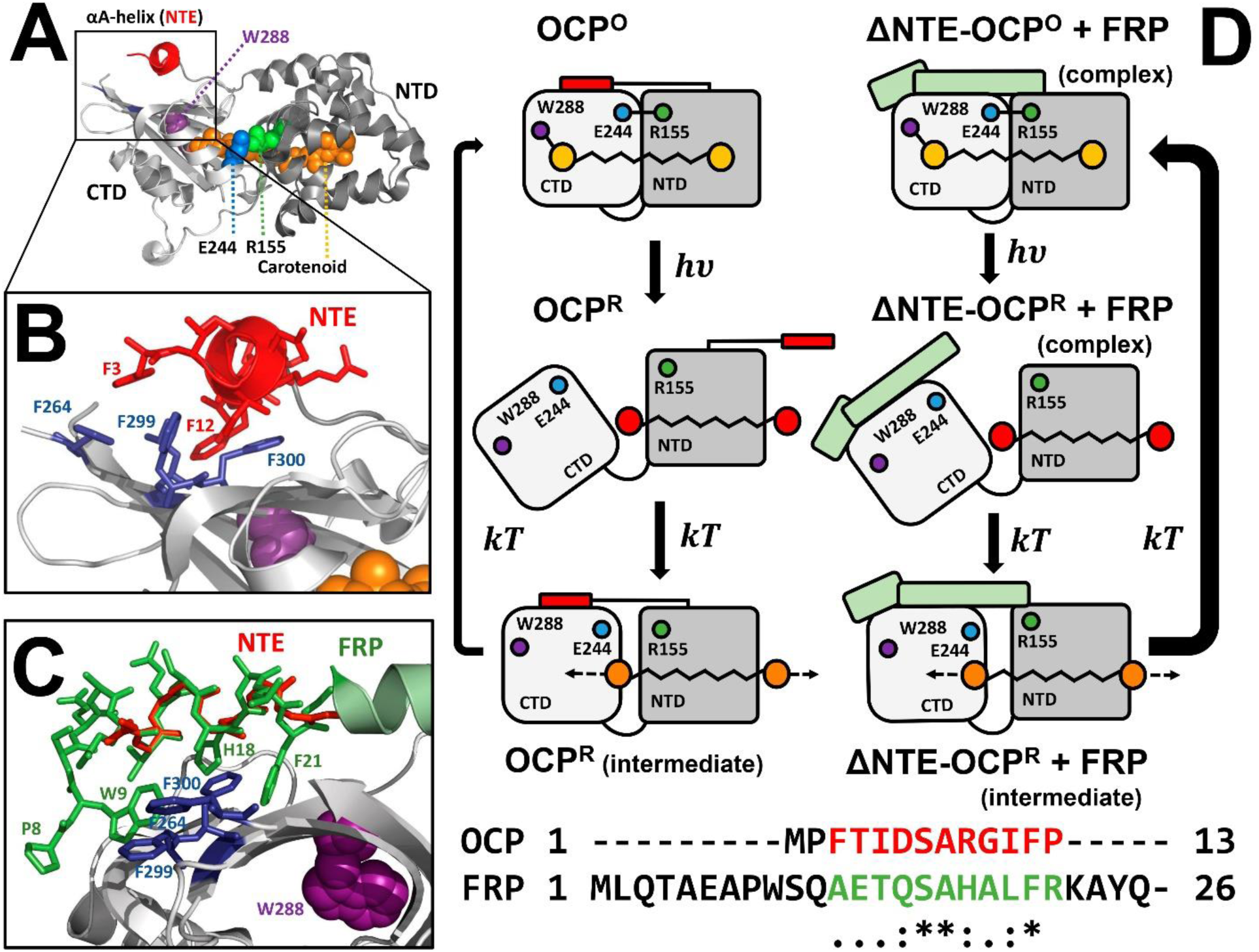
Structural and kinetic model of FRP competition with the NTE of OCP for the binding site on the OCP-CTD to expedite R→O conversion. (**A**) The OCP structure in the orange form is stabilized due to multiple chromophore/protein and protein/protein interactions: i) hydrogen bonds between the keto group of carotenoid and Trp-288/Tyr-201, ii) the Arg-155/Glu-244 salt bridge and iii) hydrophobic interactions between the residues from αA-helix of the NTE and CTD (**B**). **C**. Superposition of the α-helical FRP and NTE structures on the basis of the alignment of primary structures of the NTE of OCP (residues 3-13) and the N-terminal segment of FRP (residues 12-21) reveals similarities and suggests that FRP may mimic the NTE to compete with it for OCP binding at the overlapping site. (**D**) Schematic representation of ΔNTE-OCP-FRP **molecular rodeo mechanism** (right) in comparison with the photocycle of wildtype OCP in the absence of FRP (left) (See text for more details).

## 4. Materials and methods

### 4.1 Protein cloning, expression and purification

Cloning, expression and purification of the His6-tagged *Synechocystis* OCP^WT^ and FRP were described previously [27]. The OCP variant devoid of the N-terminal extension (NTE), termed ΔNTE herein, was constructed by recombinant PCR using a forward adapter primer introducing a *BamHI* restriction site before the codon of amino acid Val-21, a standard pQErev reverse primer, and the cDNA of OCP^WT^ as a template. Subsequent subcloning of the BamHI/NotI-restricted PCR fragment into the pQE81L vector resulted in the N-terminal amino acid sequence: MRGSHHHHHHTDPV(21)… (*Synechocystis* OCP numbering), as verified by DNA sequencing.

Holoforms of OCP^WT^ and ΔNTE were expressed in echinenone (ECN) and canthaxanthin (CAN)-producing *Escherichia coli* cells essentially as described before [26-28, 30]. All proteins were purified by immobilized metal-affinity and size-exclusion chromatography to electrophoretic homogeneity and stored at +4 °C in the presence of sodium azide. The holo-ΔNTE(O) form, separated from the ΔNTE(P) form and the apoprotein, gave the characteristic Vis/UV ratio of ~1.7.

### 4.2 Absorption spectroscopy

Absorption spectra were recorded using a Maya2000 Pro (Ocean Optics, USA) spectrometer as described in [28, 30]. Upon absorption measurements, a blue light-emitting diode (LED) (M455L3, Thorlabs, USA), with a maximum emission at 455 nm was used for the photoconversion of the samples (further - actinic light (AL) for OCP°→OCP^R^ photoconversion). Temperature of the sample was stabilized by a Peltier-controlled cuvette holder Qpod 2e (Quantum Northwest, USA) with a magnetic stirrer.

### 4.3. Analytical size-exclusion chromatography

To study concentration dependences of hydrodynamics of proteins and the interaction between FRP and ΔNTE, we pre-incubated protein samples (100 μL) and subjected them to size-exclusion chromatography (SEC) on a calibrated Superdex 200 Increase 10/300 column (GE Healthcare) equilibrated with a 20 mM Tris-HCl buffer, pH 7.6, containing 150 mM NaCl, 0.1 mM EDTA, and 3 mM Pβ-mercaptoethanol and operated at 25 °C at 1.2 mL/min flow rate. Unless otherwise indicated, the elution profiles were followed by carotenoid-specific absorbance (wavelengths are specified). In some cases, the column was constantly illuminated by a blue-LED to achieve OCP photoconversion. The binding curves were approximated using a quadratic equation allowing for simultaneous determination of apparent affinity and stoichiometry [28]. All experiments were performed at least two times using independent preparations of proteins.

## 5. Chemical crosslinking

Pre-incubated samples containing individual FRP (17.1 μM), the orange variant of ΔNTE, ΔNTE(O) (8.5 μM), OCP^WT^ (8.1 μM), or the mixtures of FRP (17.1 μM) and either ΔNTE(O) (8.5 μM) or OCP^WT^ (8.1 μM) were mixed with fresh glutaraldehyde (GA) up to a final concentration of 0.05 % for 15 min at 30 °C and then analyzed by SDS-PAGE. Control samples were mixed with buffer (10 mM HEPES, pH 7.6) lacking GA.

---

**Figure S1.**
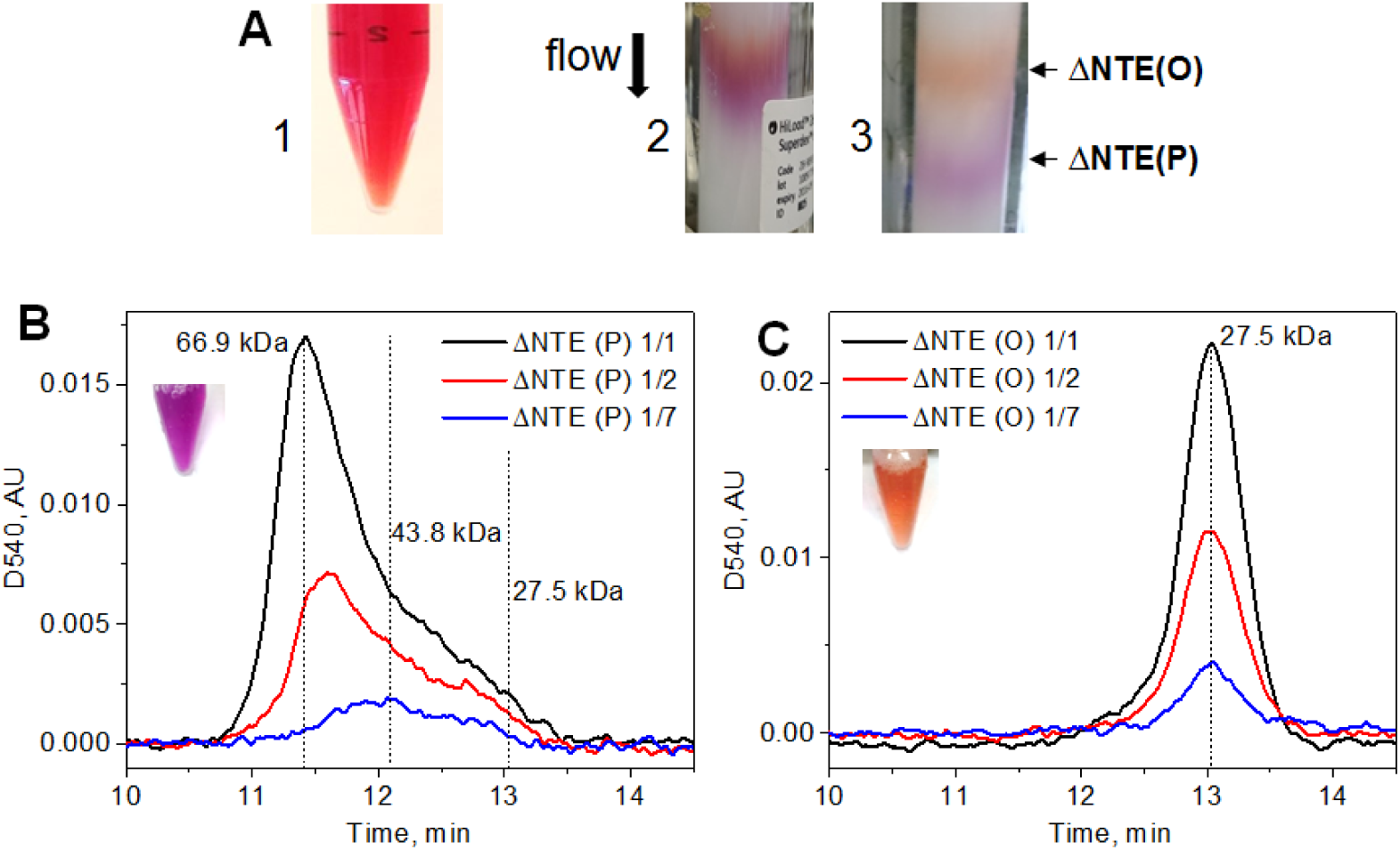
Purification and initial characterization of the ΔNTE variant of OCP. (A) The affinity purified protein (1) was separated on preparative SEC (2, 3) into two fractions, ΔNTE(P) and ΔNTE(O) with distinct properties. Panels (**B**) and (**C**) show the effect of protein concentration in the loading sample on the elution of ΔNTE(P) (**B**) and ΔNTE(O) (**C**) from the analytical Superdex 200 Increase 10/300 column (GE Healthcare). The flow rate was 1.2 mL/min.

**Figure S2.**
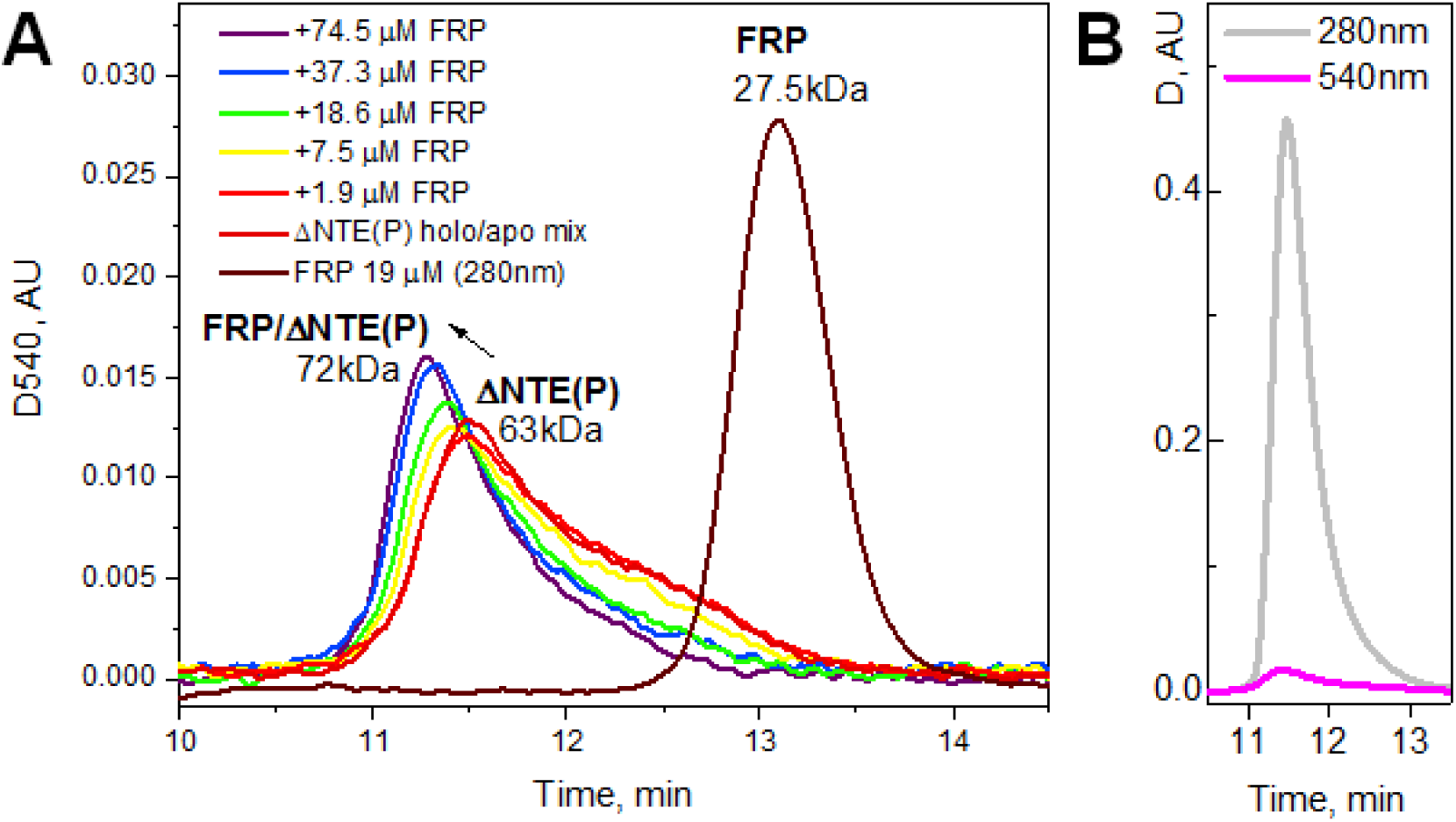
Interaction of the ΔNTE(P) sample with FRP studied by SEC. (**A**) SEC profiles for ΔNTE(P), FRP, or their mixtures at increasing FRP concentration (in the range 0-74.5 μ M) obtained using a Superdex 200 Increase 10/300 column at a flow rate of 1.2 mL/min. Positions of ΔNTE(P), FRP, and their complex as well as their apparent molecular weights are indicated. (**B**) The SEC profiles for ΔNTE(P) monitored by absorbance at two wavelengths showing that this sample represents a mixture with a very high amount of apoprotein, which complicates quantitative analysis of the ΔNTE(P)/FRP interaction.

## Acknowledgements

This work was supported by grants from Russian Foundation for Basic Research 15-04-01930a to E.G.M. E.G.M. was supported by Dynasty Foundation Fellowship. The reported study was funded by RFBR and Moscow city Government according to the research project № 15-34-70007 «mol_a_mos». T.F. acknowledges the support by the German Federal Ministry of Education and Research (WTZ-RUS grant 01DJ15007) and the German Research Foundation (Cluster of Excellence “Unifying Concepts in Catalysis”).

## Author contributions

NNS – designed and performed experiments, analysed data and wrote the paper, YBS – performed experiments, MM – performed experiments, TF – designed experiments and wrote the paper, EGM – designed and performed experiments, analysed data and wrote the paper.

